# A somatic piRNA pathway regulates epithelial-to-mesenchymal transition of chick neural crest cells

**DOI:** 10.1101/2021.04.30.442165

**Authors:** Riley Galton, Katalin Fejes-Toth, Marianne E. Bronner

**Affiliations:** Division of Biology and Biological Engineering, California Institute of Technology, Pasadena, CA 91125

**Author notes:** co-senior authors (Katalin Fejes-Toth, Marianne E. Bronner).

## Abstract

In the metazoan germline, Piwi proteins play an essential regulatory role in maintenance of stemness and self-renewal by piRNA-mediated repression of transposable elements. To date, the activity of Piwi proteins and the piRNA pathway in vertebrates was believed to be confined to the gonads. Our results reveal expression of Piwil1 in a vertebrate somatic cell type, the neural crest–a migratory embryonic stem cell population. We show that Piwil1 is expressed at low levels throughout chick neural crest development, peaking just before neural crest cells undergo an epithelial-to-mesenchymal transition to leave the neural tube and migrate into the periphery. Importantly, loss of Piwil1 impedes neural crest emigration. Small RNA sequencing reveals somatic piRNAs with sequence signatures of an active ping pong loop. Coupled with Piwil1 knockout RNA-seq, our data suggest that Piwil1 regulates expression of the transposon derived gene ERNI in the chick dorsal neural tube, which in turn suppresses Sox2 expression to precisely control the timing of neural crest specification and emigration. Our work provides mechanistic insight into a novel function of the piRNA pathway as a regulator of somatic development in vertebrates.

## Intro

Small non-coding RNAs and their protein partners, Argonaute proteins, play central regulatory roles in transcriptional and post-transcriptional gene expression in all domains of life^1^. The Piwi clade of Argonaute proteins is unique to metazoa, where they are required for maintenance of stemness, selfrenewal and safeguarding of the genome by repressing transposable elements (TEs) in the germ cell lineage^2–4^. TEs are mobile genetic elements that can replicate and reinsert themselves in the genome, threatening genomic integrity. By keeping these “selfish genes” in check, the piRNA pathway helps preserve genomic stablilty and thus plays a critical role in the arms race between TEs and their host genomes^5^. Piwi-interacting RNAs (piRNAs) recognize TE transcripts via sequence complementarity, which in the cytoplasm leads to TE target cleavage by the Piwi protein. In several organisms, Piwi proteins have gained the ability to enter the nucleus and instigate chromatin modifications at target loci. piRNA biogenesis differs from that of their better-known relatives, miRNAs and siRNAs, and relies in part on cleavage by Piwi proteins themselves in an amplification cycle termed the ping-pong amplification loop^3,6^, as well as on other cytoplasmic factors that are unique to this pathway^7,8^. These differences lead to characteristics of piRNAs that are distinct from other small RNAs: piRNAs are slightly longer than miRNAs at around 23-30 nucleotides (nt), predominantly map to TE sequences, have a 1U bias at their 5’ end and, in the case of ping-pong generated piRNAs, a 10A bias and a characteristic 10nt overlap between complementary sequences^3,6,9^. Due to 3’ end processing they also carry a 2’ O-methyl residue, which enables differential cloning^10^.

TEs are extremely prevalent and found in all metazoan genomes. New lineage-specific transposable elements emerge often during the course of evolution, and the repertoire of active transposons can vary widely among species^11,12^. Despite their negative impact on genomic stability when left unchecked, TEs provide a steady source of germline mutations and are considered an important driver of evolution. Interestingly, there are many examples of host genome cooption of retroviral genes, as well as domestication of long terminal repeats (LTRs) that bind host transcription factors to rewire gene regulatory networks, many of which act in somatic tissues^11,13,14^.

While Piwi proteins and the piRNA pathway perform critical functions in the germline to repress TEs, their potential role in somatic cells is not as well understood. It has long been queried whether the piRNA pathway might be active outside of the germline or utilized to regulate genes other than TEs. In invertebrate models like Hydra and Planaria with high regenerative capacity, somatic stem cells have been shown to utilize the canonical piRNA pathway to actively repress TEs^15,16^. In addition, Planaria Piwi protein SMEDWI-3 can regulate non-transposon mRNAs^17^. In the sea slug *Aplysia*, a piRNA mediated mechanism has been shown to mediate epigenetic regulation of synaptic plasticity^18^. piRNAs and piRNA pathway genes are also expressed in somatic tissues of numerous arthropods^19^. These findings suggests that somatic piRNA-mediated regulation may be widespread. In contrast to these invertebrate models, a functional role for the piRNA pathway in somatic cells of vertebrates has been debated^20,21^. Piwi protein expression has been observed in cancer cells^22^ and some adult tissues including hematopoietic stem-cells^23^, brain and heart^24,25^. Loss of mouse Piwi protein Mili has been correlated with behavioral deficits^25^, and Piwil1 has been implicated in neuronal migration in rats^26^, though mechanistic insights remain elusive and whether or not there is piRNA involvement in these processes is unclear.

Here we report somatic Piwi and piRNA expression in a vertebrate embryonic cell type, the avian neural crest. Our results reveal spatiotemporally regulated chick Piwil1 (Chiwi) expression and an active somatic piRNA pathway during neural crest development. The neural crest is a rapidly-evolving, migratory population of stem cells that is unique to vertebrates and essential for their development and evolution^27^. While neural crest cells undergo specification within the forming central nervous system during neurulation, they subsequently leave the neural tube to migrate extensively throughout the embryo and contribute to a diverse array of tissues^28,29^. Our functional analysis shows that Chiwi is required for neural crest cell emigration from the neural tube. Importantly, Chiwi regulates expression of a TE derived gene, ERNI, a regulator of Sox2 expression during early nervous system formation. Our results indicate that the gene regulatory network controlling chick neural crest development has coopted a transposon-derived sequence as well as its piRNA-mediated regulation to precisely time neural crest specification and initiation of the epithelial-to-mesenchymal transition (EMT) from the neural tube to begin neural crest migration.

## Results

### Chiwi exhibits a distinct expression pattern in the developing neural tube

As a first step in examining whether the piRNA pathway plays a role in regulating vertebrate developmental events, we assessed the expression of piwi genes at early embryonic stages. Vertebrates have two conserved Piwi proteins, Piwil1 and Piwil2, which play central roles in the germline piRNA pathway^30–32^. To test if *Piwi* transcripts are expressed in somatic cells of the early vertebrate embryo, we examined *Piwil1* (*Chiwi*) and *Piwil2* (*Chili*) expression during chick embryogenesis by quantitative RT-PCR using whole embryo cDNA from stages ranging from Hamburger and Hamilton (HH) stages^33^ HH4 to HH23 (Fig 1A). We observed extremely low *Chili* transcript levels at all stages. In contrast, *Chiwi* transcript levels were markedly higher, especially at early stages, peaking at HH6-8, corresponding to neurulation stages, and dropping after HH10-12, corresponding to the time of active cranial neural crest migration. Primordial germ cells are extraembryonic until stage HH13 (~2 days of development), when they enter the embryonic bloodstream through which they migrate to the gonadal anlagen by around HH18 (3 days)^34^. Thus, high *Chiwi* mRNA levels prior to this embryonic stage indicate that *Chiwi* is expressed in somatic cells.

**Figure1:**
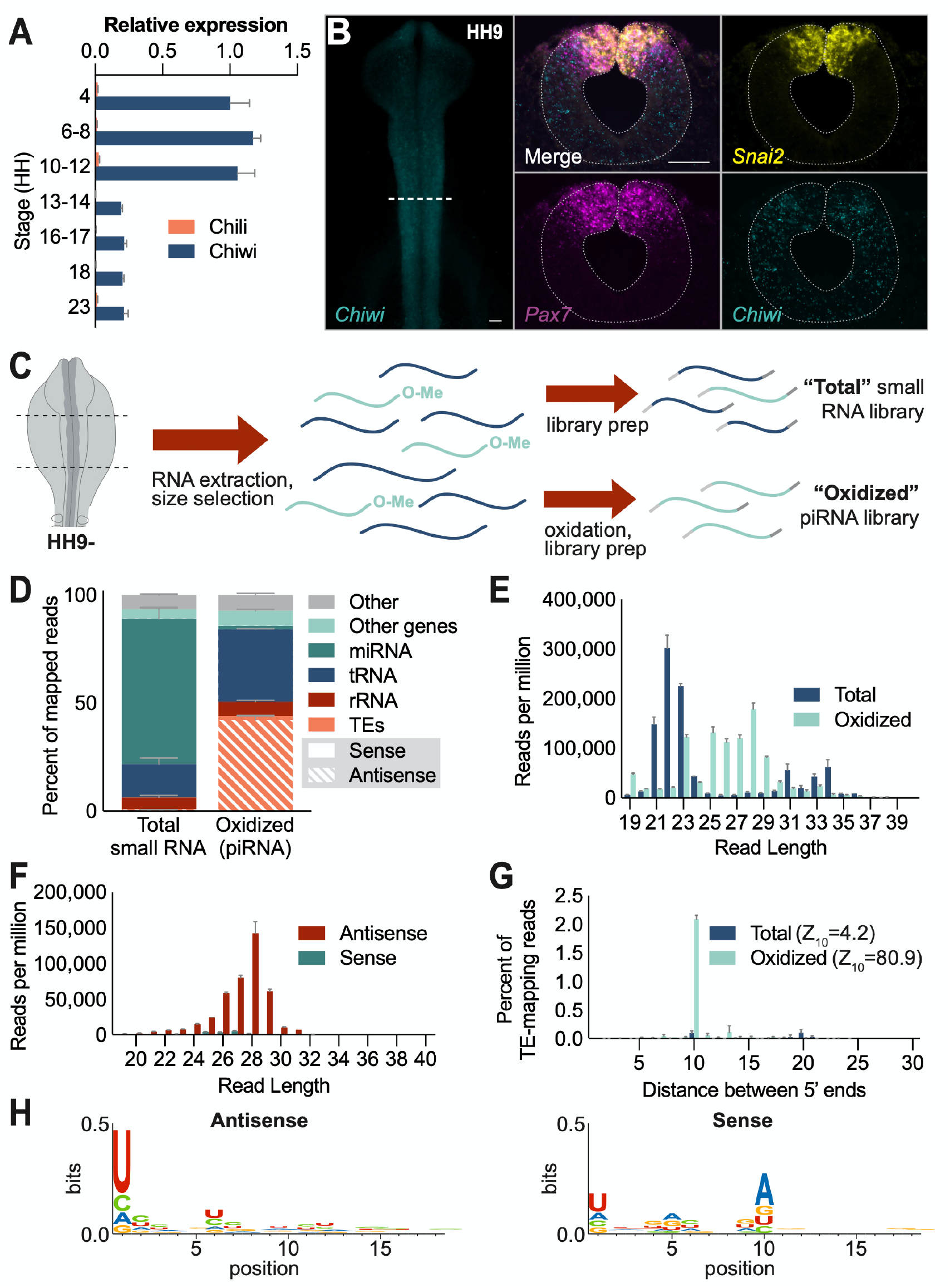
Piwi protein and piRNA expression in the cranial region. A) RT-qPCR from whole embryo RNA across stages HH4-HH23 of chick development depicts relative expression of chick piwi transcripts *Chiwi* (Piwil1) and *Chili* (Piwil2) normalized to 18S rRNA. Error bars indicate st. dev., n=1 biological replicate (RNA extracted from 3 or more pooled embryos), 3 technical replicates. B) HCR reveals expression of *Chiwi* and neural crest markers *Pax7* and *Snai2*; scale bars = 50μm. C) Schematic diagram of the small RNA cloning strategy from the midbrain region of HH9-embryos (n=2 biological replicates). D) Annotation of small RNAs mapping to the genome. Orientation is relative to the annotated feature. “Other” category includes reads that could not be assigned to a feature, as well as reads mapping to simple repeats, satellite repeats, scRNA and snRNA, which together account for <1% of mapped reads in all samples. E) Length distribution of all reads mapping to the genome in total and oxidized libraries. F) Length distribution of reads from oxidized libraries mapping to TEs in sense and antisense orientation. G) Analysis of 5’ to 5’ distance of complementary small RNA reads mapping to TEs in total and oxidized libraries. H) Sequence logos of oxidized, collapsed sequences mapping to TEs in antisense (left) and sense (right) orientation.

To confirm the RT-qPCR results and assess *Chiwi* expression with spatiotemporal resolution, we performed hybridization chain reaction (HCR) on HH6-12 embryos in whole mount (Fig 1B, SupFig 1A). Transverse sections revealed low but ubiquitous Chiwi expression in the cranial region at HH6, which becomes primarily constrained to the neural tube from HH9 and onward (Fig 1B, SupFig 1B). Interestingly, Chiwi expression appears to be differentially regulated in the dorsal tube at HH9, where there are distinguishable subdomains of neural crest precursors that differ in their expression of neural crest specifier genes^28,35^ (Fig 1B). In the dorsal midline, there are cells that express both *Snai2* and *Pax7*, corresponding to premigratory neural crest cells in the process of undergoing EMT^28,36^. In this domain, *Chiwi* expression is reduced compared to the rest of the neural tube. In contrast, in the immediately lateral domain which feeds into the specified neural crest pool^28^, cells are marked by high *Pax7* but no *Snai2* and relatively high levels of *Chiwi* expression.

We next confirmed the presence of *Chiwi* in the neural crest region by RNA-seq. To this end, we used three RNA-seq datasets, one from dissected cranial neural folds (two replicates) and two previously published FACS sorted specified neural crest datasets^37^, which include early migrating cranial neural crest cells from HH9 stage embryos (three replicates), and trunk neural crest cells from HH18 embryos (three replicates). All three datasets reveal notable expression levels of *Chiwi* mRNA, with a respective transcripts per million (TPM) of 2.8 and 2.6 in the specified cranial and trunk datasets, and a TPM of 6.5 in the cranial neural fold dataset. Interestingly, Chili was detected in neural folds with a TPM of 4.1 but was not detectable in sorted migrating neural crest cells (SupFig 1C). Taken together, these results support the somatic expression of *Chiwi* in both neural crest precursors and early migrating neural crest cells of the chicken embryo, and low-level expression of *Chili* in dorsal neural folds.

### Somatic piRNAs target transposons

To address whether Chiwi could exert a regulatory function in the neural crest that is directed by associated piRNAs, we tested for the presence of piRNAs in the cranial region. Due to the lack of appropriate antibodies, it was not feasible to immunoprecipitate Chiwi and directly analyze associated small RNAs. As an alternative, we cloned and sequenced small RNAs from the midbrain region of HH9 heads, where *Chiwi* expression is very pronounced and confined to the neural tube. In parallel, to specifically enrich for piRNAs, we performed small RNA cloning that included an oxidation step to select for RNA species that are 2’-O-methylated at their 3’ end, a characteristic of piRNAs (Fig 1C).

Roughly 64% and 60% of the reads in the total and oxidized samples mapped to the genome with no mismatches. Amongst these, both the total and oxidized samples showed comparable numbers of reads mapping to rRNA, at around 6% (Fig 1D). Total small RNA reads mostly mapped to miRNAs, with some mapping to tRNAs and to genes in sense orientation (Fig 1D). Sense mapping reads are unable to target the corresponding mRNAs and likely represent degradation products. Less than one percent of mapped reads in the total small RNA samples corresponded to transposon sequences, mostly in antisense orientation, confirming that piRNA expression in this tissue is extremely low relative to other small RNA species. The representation of different TE families in total TE mapping reads, however, was similar to the oxidized samples, indicating that the total small RNA library contains piRNAs, even if they represent only a small fraction of all reads (SupFig 2A). Consistent with predominance of miRNAs and degradation products in the non-oxidized libraries, the size distribution of the total small RNA samples peaked at 22 nucleotides and showed a broader distribution across all sizes (Fig 1E). In contrast, 44% of the reads in the oxidized samples mapped to TEs with 96% of these in antisense orientation (Fig 1D), and the size distribution of these libraries peaked at 28 nucleotides (Fig 1E), consistent with the size profile of previously published chick embryonic piRNA libraries^38^. Interestingly, we noted an enrichment of tRNA mapping reads in addition to TE mapping reads in the oxidized libraries, consistent with tRNAs being heavily 2’O methylated^39^ (Fig 1D). Upon plotting the size distribution of tRNA mapping reads, we saw a strong enrichment of 23 and 25nt sequences in the oxidized samples, possibly suggesting the presence of tRNA-derived small RNAs with 2’-O-methylated 3’ ends (SupFig 2B).

To further characterize the presumptive piRNA population, we analyzed the size profiles of sense and antisense TE-mapping reads in the oxidized samples. Interestingly, we found that sense reads peaked between 25-27 nucleotides, while the more abundant antisense reads had a strong peak at 28 nucleotides (SupFig 2C, Fig 1F). While the RT-qPCR and RNA-seq data indicated very low Chili expression (Fig 1A, Supfig 1C), these size profiles suggest that more than one Piwi protein might be present —albeit at different expression levels— and contributing to piRNA biogenesis via the ping-pong amplification mechanism. Consistent with this idea, we observed a strong 10 base pair overlap between complementary reads in the oxidized libraries when we analyzed 5’ to 5’ overlap of TE-mapping reads (Fig 1G), as well as a notable 1U bias of antisense and 10A bias of sense sequences upon collapsing the libraries (Fig 1H). These findings provide strong support for an active somatic piRNA pathway in the embryo, and imply that somatic piRNAs are, at least in part, generated by a ping-pong mechanism, possibly through the interplay of two Piwi proteins.

To determine if somatic piRNAs might be generated by a similar mechanism to germline piRNAs, we looked for expression of genes involved in germline piRNA biogenesis in RNA-seq data from dorsal neural folds (SupFig 3). Interestingly, several key factors are expressed at low levels, comparable to those of *Chiwi* and *Chili*. These include Hen1 (TPM 3.3) which imparts 2’-O-methylation to piRNAs^7^, TDRD9 (TPM = 0.6) which is involved in ping-pong piRNA biogenesis^7^, PNLDC1 (TPM 4.3) which trims the 3’ ends of piRNAs^8^, and possibly the Zucchini homolog PLD6 (TPM = 13.1)^7^, though RNA-seq tracks of PLD6 are noisier than those of the other genes. Interestingly, the transcription factor A-MYB, which mediates expression of both piRNA clusters and piRNA pathway genes, is also present (TPM = 1.4), as are the the two tudor domain containing genes TDRD1 (TPM=3.9) and TDRD3 (TPM=3.6) which A-MYB has been shown to regulate along with Chiwi in rooster testes^40^. Altogether, expression of piRNA pathway genes in the dorsal neural tube suggests a similar mechanism of piRNA biogenesis to that seen in the germline.

### Chiwi is required for proper neural crest development

Based on Chiwi’s distinct expression pattern, we hypothesized that it plays a role in neural crest development. To probe Chiwi’s function in the neural crest, we first performed loss of function experiments using a translation blocking Chiwi morpholino oligomer (MO). After electroporating Chiwi MO into the prospective neural crest region on one side of the embryo and control MO on the other side at HH4, we allowed the embryos to grow until HH9 when cranial neural crest begins to delaminate and migrate and Chiwi appears to be relatively strongly expressed in the dorsal neural tube (Fig 2A). We observed a reduction in neural crest migration distance from the midline on the Chiwi depleted side by immunostaining for the neural crest marker Pax7. Upon sectioning, we saw significant reduction in the numbers of Pax7 positive cells on the Chiwi depleted side (Fig 2B).

**Figure 2:**
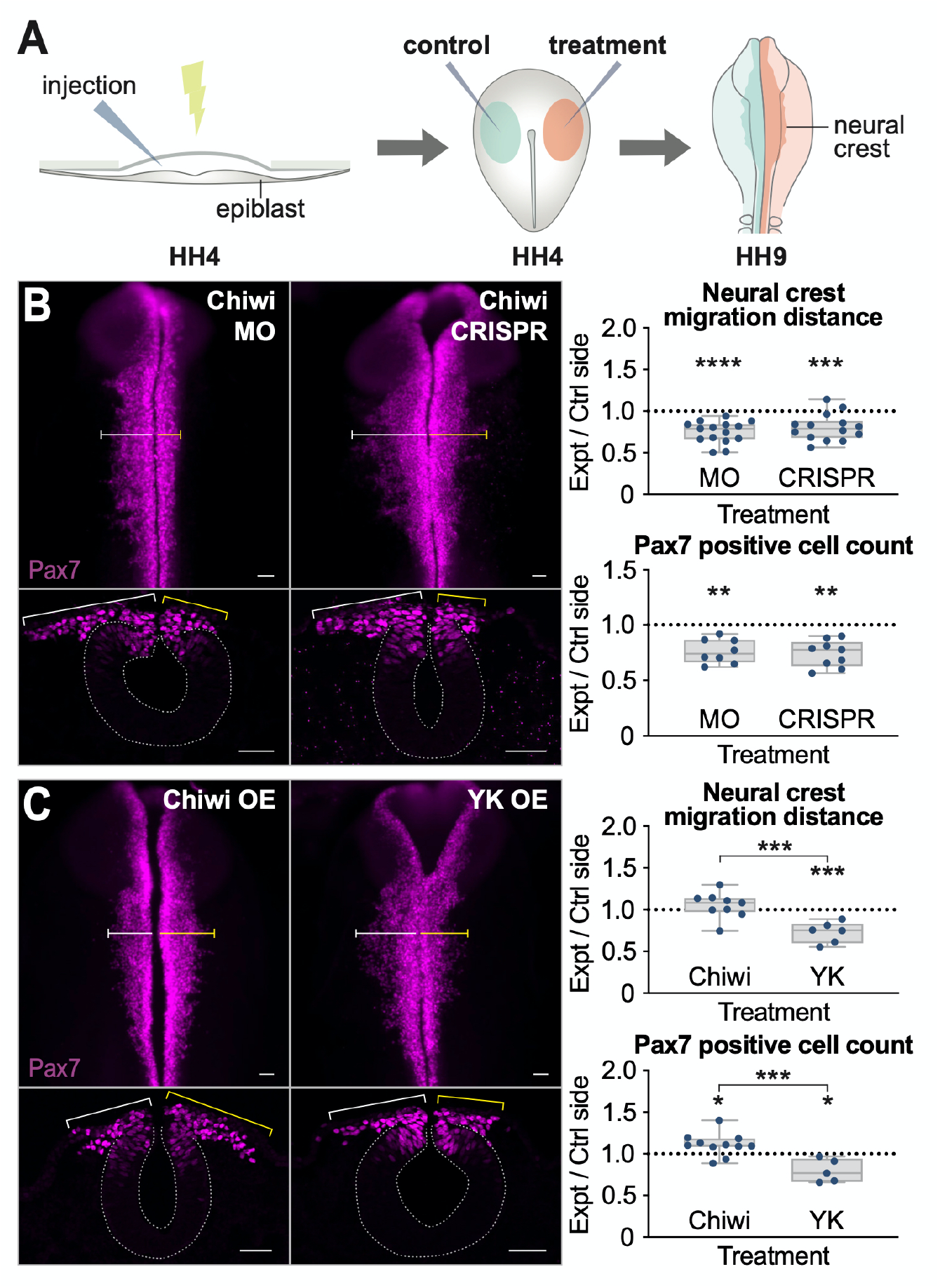
Chiwi is required for neural crest development. A) Schematic diagram of the chick embryo electroporation strategy. B) Loss of Chiwi impedes neural crest migration and reduces neural crest cell count, as measured by Pax7 expressing cells. Left: examples of whole mount and cross sections upon morpholino (MO) and CRISPR knockout of Chiwi. Right: quantification of image analysis. Each data point represents measurements from a single embryo, with the right (experimental) divided by the left (control) side. C) Overexpression (OE) of Chiwi increases the number of Pax7 positives cells, though migration distance is not significantly altered, while overexpression of the YK mutant, which is unable to bind piRNAs, impedes neural crest migration and reduces cell number. All scale bars = 50μm. Box plots indicate the interquartile range, while whiskers extend to min and max values. *, **, *** and **** indicate p values of ≤ 0.05, 0.01, 0.001, and 0.0001, respectively, and represent the difference between control and experimental meausrements for each treatment.

To confirm the specificity of our results, we used a second approach to knock-out *Chiwi* via a plasmid based CRISPR/Cas9 system optimized for use in chicken^41^. To this end, we designed two CRISPR guide RNAs against sequences corresponding to the first exon junction and the 3’ piRNA binding site of Chiwi. We electroporated a Cas9 construct along with control guide on one side and *Chiwi* guides on the other side of HH4 stage embryos. HCR for *Chiwi* after CRISPR/Cas9 knockout confirmed a reduction in *Chiwi* mRNA levels compared to the control side at stage HH9, just prior to onset of neural crest migration (SupFig 4). CRISPR-Cas9 mediated loss of *Chiwi* gave the same reduction in neural crest migration distance and cell count at HH9 as did the Chiwi morpholino (Fig 2B).

### Chiwi’s piRNA-binding activity is required for its function in the neural crest

Based on the presence of piRNAs in the cranial midbrain region, we hypothesized that Chiwi identifies its targets via the associated piRNAs. To test the need for Chiwi’s piRNA binding activity, we created wild type and mutant Chiwi overexpression constructs. The mCMV-YK-Chiwi construct contains two amino acid substitution in positions 574 and 578, resulting in impaired piRNA binding^42^. To express these at levels similar to those of endogenous Chiwi, cDNA amplicons were cloned into a construct containing a minimal CMV (mCMV) promoter, which has weak expression in chick. The vector also contains an H2B-RFP sequence following an internal ribosome enry site (IRES) downstream of the Chiwi sequence. As a control, we employed the empty vector, which expresses only the H2B-RFP under the mCMV promoter. While overexpression of wild type Chiwi did not significantly alter neural crest migration, it did result in an increase in Pax7 positive cells. In contrast, overexpression of the piRNA binding mutant resulted in reduced neural crest migration distance and cell count (Fig 2C) compared to the control, recapitulating Chiwi loss of function and indicating that Chiwi’s piRNA binding ability is required for its function in the neural crest.

### Chiwi regulates a single, transposon-derived gene, ERNI in the dorsal neural tube

To investigate Chiwi targets in the neural crest, we performed RNA-seq experiments after unilateral CRISPR-Cas9 mediated knock-out. Differential mRNA expression analysis of control and knock-out RNA-seq libraries generated from total RNA of neural folds at stage HH9 revealed that the most significantly upregulated gene upon Chiwi knockout is the retrotransposon ENS-1, also called Soprano (Fig 3A). ENS-1 has numerous copies in the genome, most of which only contain the LTR^43,44^. Importantly, several copies contain only the 5’ end of the internal domain, which codes for ERNI^43,45^, a chicken-specific transposon-derived gene and known regulator of neural induction^46,47^.

**Figure 3:**
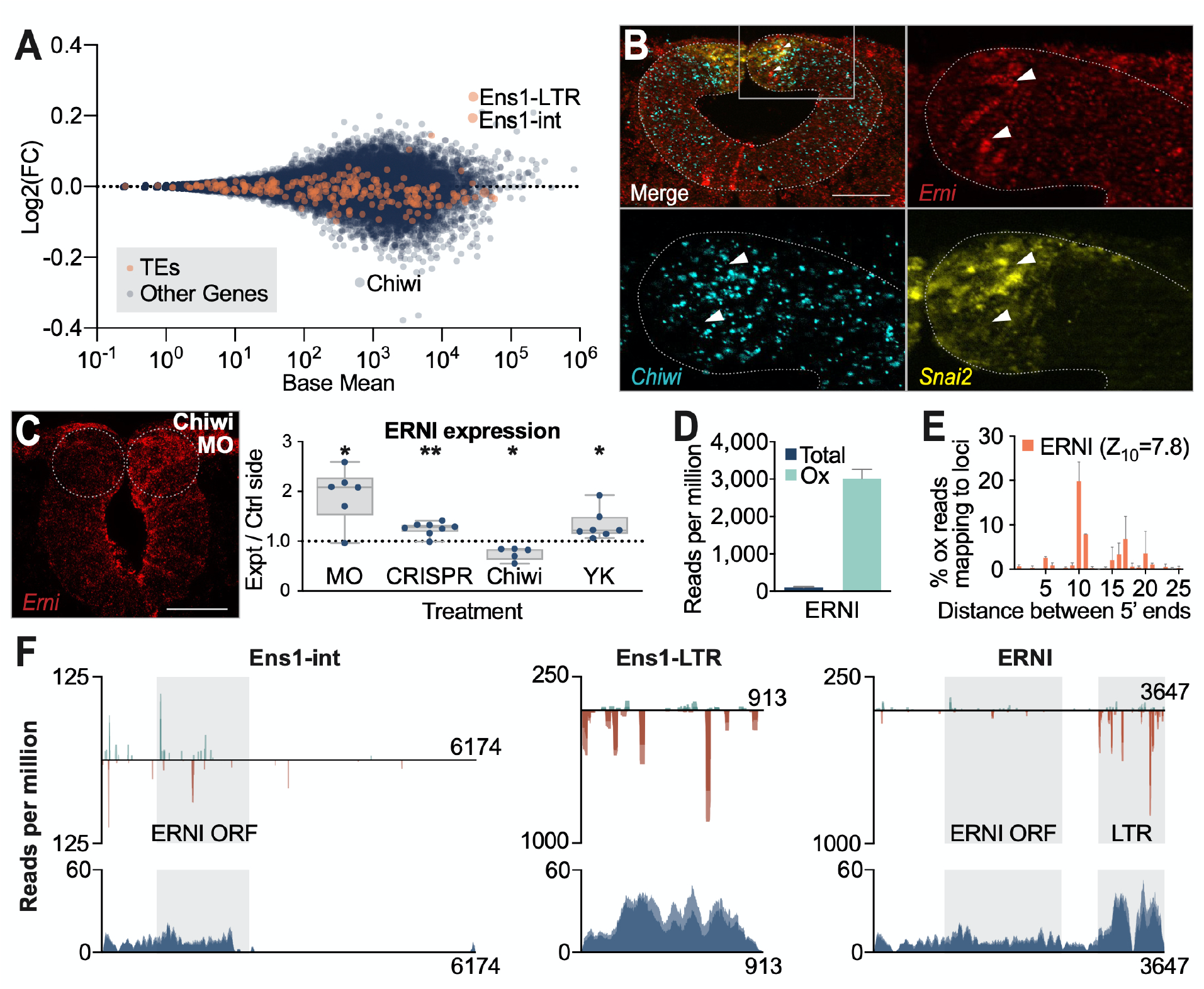
Chiwi regulates a single, transposon-derived gene, ERNI, in the neural crest. A) Differential expression analysis of RNA-seq from HH9-Chiwi CRISPR knockout vs control cranial neural folds. Dots represent protein coding genes (blue) and transposon families (orange). B) HCR depicting *ERNI*, *Chiwi* and *Snai2* expression in WT HH9 cranial midbrain region; scale bar = 50μm. Arrowheads point to two ERNI positive cells for spatial comparison of gene expression. C) HCR reveals changes to *ERNI* expression upon Chiwi loss (MO and CRISPR), as well as Chiwi and YK mutant overexpression. Each data point represents the average fluorescent intensity of the right dorsal fold region (experimental) divided by the left (control) side from three non-adjacent sections from the cranial region of a single embryo. Scale bar = 50μm. Box plots indicate the interquartile range, while whiskers extend to min and max values. *, **, *** and **** indicate p values of ≤ 0.05, 0.01, 0.001, and 0.0001, respectively, and represent the difference between control and experimental measurements for each treatment. D) Normalized small RNA read counts mapping to the ERNI mRNA sequence in total vs oxidized small RNA libraries. Error bars indicate st. dev. from two biological replicates. E) Analysis of 5’ to 5’ distance of complementary small RNA sequences mapping to the ERNI mRNA sequence in the oxidized small RNA libraries. F) Small RNA-seq (top) and RNA-seq (bottom) tracks depicting sequences mapping to ENS1 loci as well as the ERNI mRNA sequence (left). Oxidized small RNAs mapping in sense (teal) and antisense (red) orientation are depicted separately. Cranial neural fold total RNA-seq (control libraries from A) are depicted in blue. Replicate tracks are overlaid.

To validate our RNA-seq results, we performed HCR against the ERNI mRNA sequence (Fig 3B). In wildtype embryos, this revealed ERNI expression throughout the neural tube and ectoderm, albeit at varying levels. In particular, we noted higher expression in the dorsal most-region of the neural tube, corresponding to the Snai2+ domain in which Chiwi transcripts are downregulated. Consistent with this, ERNI was reduced compared to the control side after Chiwi overexpression, whereas Chiwi depletion and overexpression of the piRNA binding mutant resulted in an increase in ERNI (Fig 3C).

Analysis of small RNA reads mapping to the ERNI mRNA sequence, which includes portions of both the ENS-1 internal domain and LTR, showed enrichment in the oxidized piRNA sample compared to total small RNAs, and displayed the 5’ to 5’ complementarity signature of the ping pong pathway (Fig 3D-E), indicating that Chiwi is regulating ERNI via a piRNA mediated mechanism. Interestingly, both the RNA-seq and the small RNA samples showed enrichment specifically over the LTR and ERNI sequence and did not map to other internal parts of the ENS-1 transposon (Fig 3F). Together, these results indicate that ERNI expression in the dorsal neural tube is spatially regulated by Chiwi in a piRNA-dependent fashion.

### Perturbation of ERNI recapitulates Chiwi phenotypes

To functionally test whether dysregulation of ERNI could account for the observed neural crest defects upon Chiwi perturbation, we directly disrupted ERNI expression in the dorsal neural tube and analyzed its effect on neural crest cell count and migration. To this end, we generated an overexpression vector, pCI-FLAG-ERNI, which encodes an N-terminally FLAG-tagged ERNI coding sequence followed by H2B-RFP separated by an IRES, all under the chicken-□-actin promoter, which displays strong expression in the chick embryo (Fig 4A). We electroporated this construct into one side of the embryo at HH4 with a control version containing just the H2B-RFP on the other side. Pax7 staining of HH9 embryos revealed a reduction in neural crest cell count and migration distance, similar to that seen after Chiwi knockout (Fig 4B); this is consistent with Chiwi regulating ERNI expression. The ERNI protein harbors an N-terminal coiled-coil domain, which is responsible for recruitment of ERNI to the Sox2 N2 enhancer, as well as a C-terminal HP1-box which recruits HP1-gamma, thereby inducing transcriptional repression^47^. To further test ERNI’s function, we created a dominant negative ERNI construct, pCI-N150-FLAG-ERNI-NLS (N150), which only contains the first 150 amino acids of ERNI. This includes the Sox2 localizing coiled-coil domain but lacks the HP1-gamma binding domain. It also contains a nuclear localization signal on the C-terminus to ensure nuclear localization. Overexpression of this dominant negative construct imparted a reciprocal phenotype to wildtype ERNI overexpression, with increased number of Pax7+ neural crest cells at HH9 (Fig 4B), recapitulating the Chiwi overexpression phenotype. Taken together, these data suggest that Chiwi’s primary function during neural crest development is to regulate ERNI expression in a spatiotemporal manner.

**Figure 4:**
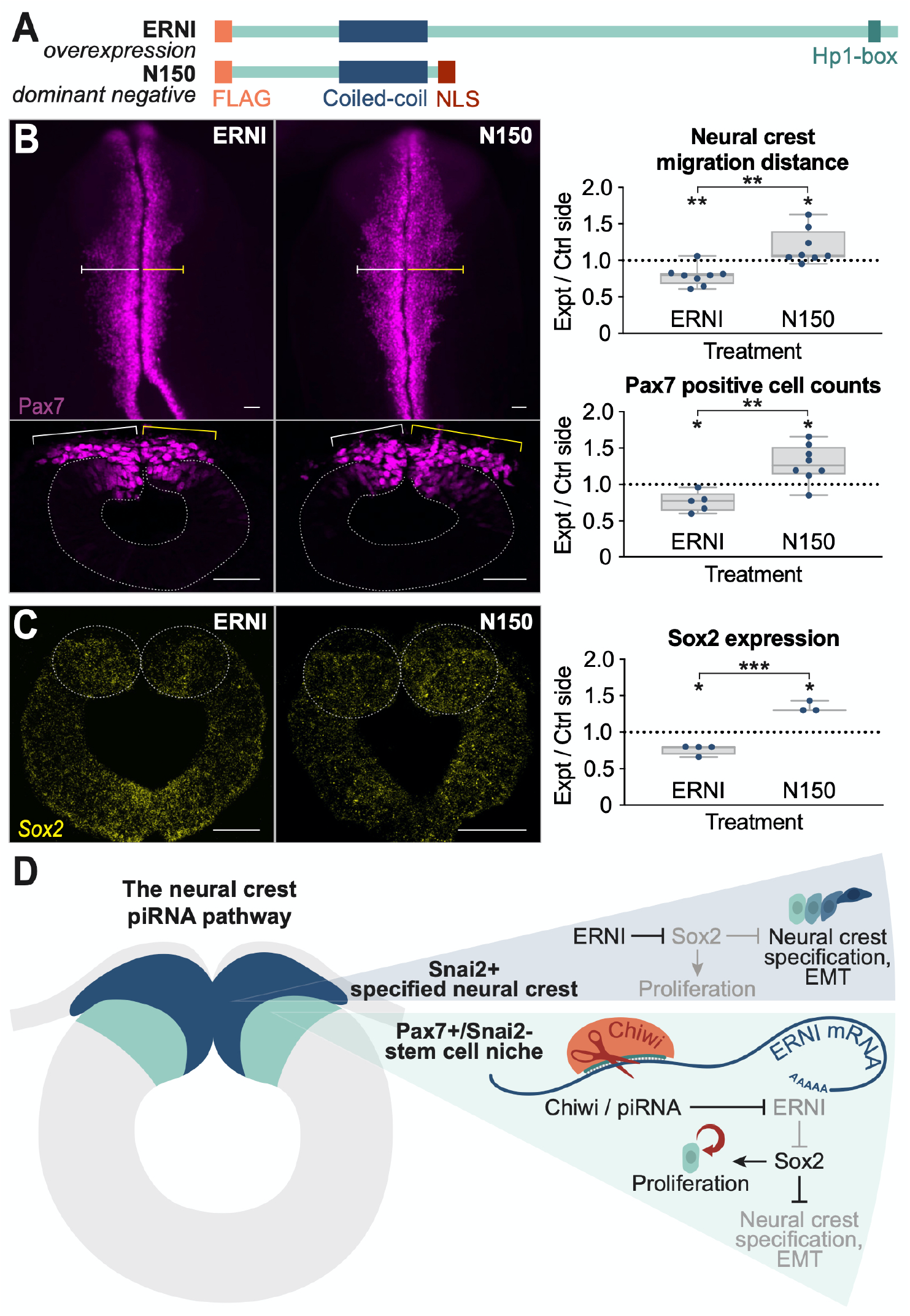
Perturbation of ERNI recapitulates Chiwi phenotypes. A) Schematic diagram of ERNI overexpression and dominant negative (N150) expression construct products. B) Overexpression of ERNI recapitulates loss of Chiwi phenotype, with a reduction in neural crest migration distance and cell number, while overexpression of the N150 truncated ERNI sequence increases cell count and migration distance, as measured by Pax7 expressing cells. Left: examples of whole mount and cross sections upon ERNI and N150 overexpression. Right: quantification of image analysis. Each data point represents measurements from a single embryo, with the right (experimental) divided by the left (control) side. C) HCR reveals that ERNI overexpression leads to a reduction in Sox2 expression in the dorsal neural tube, while N150 dominant negative overexpression increases Sox2 expression. Each data point represents the average fluorescent intensity of Sox2 signal in the right dorsal fold region (experimental) divided by the left (control) side from three non-adjacent sections from the cranial region of a single embryo. All scale bars = 50μm. Box plots indicate the interquartile range, while whiskers extend to min and max values. *, **, *** and **** indicate p values of ≤ 0.05, 0.01, 0.001, and 0.0001, respectively, and represent the difference between control and experimental meausrements for each treatment. D) Schematic diagram of the neural crest piRNA pathway, depicting a neural tube with the PAX7+/Snai2-stem cell niche in teal, which feeds into the Snai2+ specified neural crest region in dark blue. Chiwi represses ERNI in the PAX7+/Snai2-stem cell niche via a piRNA mediated mechanism in order to maintain Sox2 expression and proliferation. In specified neural crest, Chiwi expression is reduced, permitting ERNI expression which in turn represses Sox2 to allow for neural crest specification and EMT.

### ERNI regulates Sox2 during neural crest specification

Since ERNI has been shown to regulate Sox2 expression at early gastrula stages^46,47^, we next tested whether it also plays a role in Sox2 regulation in the dorsal neural tube during neural crest development. To address this, we performed HCR for Sox2 on embryos into which we electroporated the ERNI overexpression and ERNI dominant negative constructs. We noted a decrease in *Sox2* expression in the *Pax7*+ domain of embryos where *ERNI* expression is increased, and an increase in *Sox2* expression in this domain of embryos that expressed the N150 dominant negative construct (Fig 4C); this is consistent with ERNI inducing Sox2 repression. Taken together, these results suggest that ERNI functions to repress *Sox2* in the dorsal neural tube, thereby enabling activation of the gene program to specify bona fide neural crest cells, which is inhibited by high levels of Sox2^28^. Chiwi, in turn, is responsible for repressing ERNI to maintain *Sox2* expression in the *Pax7*+*/Snai2-* dorsal neural tube cells, prior to the onset of specification and emigration from the neural tube (Fig 4D).

## Discussion

Host cooption of transposable element components is a relatively common occurrence and a significant driver of evolution. Until now, the piRNA pathway had not been implicated in this process. Here we show that the piRNA pathway, canonically functioning in the germline to repress deleterious expression of TEs, has been coopted by the chicken embryo to spatiotemporally regulate the expression of TE derived gene ERNI, and precisely time neural crest epithelial-to-mesenchymal transition. To our knowledge, this is the first demonstration of cooption of the piRNA pathway to regulate a developmentally relevant gene, expanding our understanding of the piRNA pathway from solely being an antagonistic force against TEs, to playing a critical role in the domestication of a TE derived gene.

ERNI is known to regulate Sox2 at the onset of neural induction in the chicken embryo^47^ and is derived from an endogenous retrovirus sequence that appears to be unique to the galliforme lineage^43^. ERNI expression has also been observed in chicken ES cells and in the embryonic gonads, where it appears to be a marker of pluripotency ^45,48^. The fact that ERNI has embedded itself in the highly conserved processes of neural crest EMT and neural induction raises the possibility that similar regulation of development and differentiation by piRNA-mediated regulation of TE sequences might be happening in other species. Intriguingly, another TE derived gene, Crestin, is expressed in Zebrafish neural crest cells, though its function is unclear^49^. We hypothesize that the piRNA pathway plays a conserved role in vertebrate somatic development, though the transposon sequences through which it exerts its function likely change more frequently, in concert with the ever-evolving TE landscape of vertebrates. By constantly updating the pool of piRNAs to match evolving TE profiles, the piRNA pathway naturally provides the needed plasticity to repress whichever TE is co-opted by the gene regulatory network in a given species or tissue. Consistent with this idea, we observed piRNAs from several TE families in the chick midbrain, suggesting that despite the fact that only ERNI is being regulated by the piRNA pathway in this context, the machinery necessary to regulate other TEs in the same spatiotemporal manner is already present.

In the dorsal region of the chick neural tube, we posit that downregulation of Sox2 by ERNI permits Pax7 positive cells to express Snai2 and undergo EMT, becoming migratory neural crest. Interestingly, however, when we overexpress ERNI we see less Sox2 in the dorsal neural tube, but fewer neural crest cells. One possible explanation for this observation is that ERNI might also regulate other genes in addition to Sox2 in the dorsal neural tube. We postulate, however, that very precise Sox2 levels are required to maintain the proliferative ability of Pax7+/Snai2-cells–effectively reflecting a neural crest stem cell niche—such that reducing Sox2 in this region leads to a loss of their ability to self-renew. This model of Sox2’s role in neural crest development mirrors its function in other cell types. Sox2 is a well known stem cell factor, required for maintenance of several developmental stem cell niches, and precise maintenance of its expression levels can have a significant effect on cell fate. For example, a recent study found Sox2 levels in the chick tail bud modulate a stem cell population driving secondary neurulation, with very low levels of Sox2 required to maintain proliferation in the stem cell niche, and overexpression of Sox2 instigating differentiation into neural epithelium and also reducing the self-proliferative properties of these cells^50^. Thus, we hypothesize that the piRNA pathway may function as a guardian of the stem cell niche that feeds into the specified neural crest region, regulating proliferation by repressing ERNI to maintain Sox2 expression, and only permitting ERNI to switch off Sox2 when it is time for a cell to undergo EMT and become bona fide migratory neural crest.

The piRNA pathway has long been thought to be confined to the germline in most organisms because expression of its components is typically not observed in other tissues, particularly in vertebrates. Unlike the germline piRNA pathway, it is worth noting that the neural crest piRNA pathway requires relatively low levels of Chiwi and its associated piRNAs to function. The fact that Chiwi can have a significant effect at these levels suggests that there may be other somatic piRNA pathways previously missed due to low expression profiles. This in turn raises the intriguing possibility that repurposing of the piRNA pathway for somatic gene regulation may be a widespread occurrence, perhaps shedding light on the myriad reports of Piwi protein expression in diverse cancers. Somatic regulation by Piwi proteins not only leads to interesting new functions of the piRNA pathway outside of the gonads, but implies that activation of the piRNA pathway, and thus its adaptation as an epigenetic tool in research and therapy, may be an attainable goal.

## Acknowledgements

We thank members of the Bronner, Fejes Toth and Aravin labs for helpful discussions. We thank Maria Ninova for advice on RNA-seq analysis, as well as for making the pingpong script available for our use. We also thank Michael Piacentino for advice with imaging analysis and providing FIJI macros for our use. We thank Qing Tang for providing us with the mini CMV promoter. We acknowledge the Caltech Millard and Muriel Jacobs Genetics and Genomics Laboratory for library prep and sequencing of our CRISPR RNA-seq experiment, and in particular thank Igor Antoshechkin for advice on data analysis, as well as ensuring that our small RNA libraries got sequenced during the COVID-19 pandemic. This work is supported by NIH grants R01GM110217 to KFT and R35NS111564 to MB. RG was supported by the NSF GRFP fellowship.

## Author contributions

RG, MB and KFT designed experiments; RG conducted all experiments and data analysis; RG, MB and KFT wrote the manuscript.

## Data availability statement

All raw sequencing data generated for this publication is available upon reasonable request from the corresponding authors, and will available through Gene Expression Omnibus upon publication. Previously published specified neural crest datasets are available from NCBI BioProject# PRJNA497902.

## Methods

### Cloning of expression vectors

The mCMV-H2B-RFP construct was generated by replacing the rabbit β-actin promoter of PCI-H2B-RFP with a minimal CMV promoter sequence using the restriction sites Spe1 and Xba1. This promoter also contains a 5xTetO site, which we left uninduced to achieve weak expression. The Chiwi coding sequence (Ensembl transcript ENSGALT00000004171.6) was PCR amplified from HH10-12 whole embryo cDNA, prepared using oligo(dT) primers and the Superscript III reverse transcriptase kit, and subsequently cloned into the mCMV-H2B-RFP vector using the restriction sites Asc1 and Xho1 to create mCMV-Chiwi. The mutant construct (mCMV-YK-Chiwi) was generated using the QuickChange Lighting Multi Site-Directed Mutagenesis kit to generate the Y574I and K578E amino acid substitutions, which have been previously described to inhibit piRNA binding^42,51^. As a marker for electroporation efficiency, pCI-H2B-RFP was co-electroporated alongside the mCMV constructs, as mCMV-H2B-RFP expression is too low to pick up post methanol dehydration of embryos.

The ERNI coding sequence (NCBI refseq NM_001080874.1) was PCR amplified from cDNA obtained from reverse transcribing HH9 head RNA using oligo(dT), and cloned with an N-terminal FLAG tag into the pCI-H2B-RFP vector using AflII and Nhe1 (pCI-FLAG-ERNI). The coding sequence for the first 150 amino acids (N150) was similarly cloned into pCI-H2B-RFP, but with an N-terminal FLAG tag and a C-terminal nuclear localization signal (pCI-N150-FLAG-ERNI-NLS).

### Electroporation

Fertilized chicken eggs were acquired from various providers, most recently Sun State Ranch (Sylmar, CA), and grown at 37°C for 18-20 hours to reach HH4-5. Ex ovo electroporations were performed on stage HH4-5 embryos as previously described. Embryos were dissected onto rings of filter paper in Ringer’s and a solution of DNA expression construct or morpholino (MO) was injected into the space between the vitelline membrane and ectoderm (Fig 3A) and electroporated into the ectoderm with 5 pulses of 5.2V for 50ms, with 100ms between each pulse. Embryos were then cultured at 37°C in thin albumen with penicillin/streptomycin until HH9. All embryos were bilaterally electroporated with the control on the left side and experimental on the right side, allowing for direct comparison. Prior to analysis, embryos were screened for electroporation efficiency and coverage by fluorescent reporter expression and/or immunostaining of tagged proteins.

Expression constructs were injected at concentrations of 1-2.5μg/ul, while MOs were used at a concentration of 0.25mM with 1.0μg/ul pCIG-GFP as carrier DNA. FITC labelled MOs used include standard control MO (5’–CCTCTTACCTCAGTTACAATTTATA–3’) and Chiwi translation blocking MO (5’-TCTGGCTCTAGCTCTTCCTGTCATG-3’) from Gene Tools.

CRISPR mediated knockout was performed as described previously^52^. A Cas9 expressing construct (pCAG-nls-hCas9-nls-eGFP) was co-electroporated alongside two guide RNA constructs targeting Chiwi—one in the first exon (5’-GGGAGGTCTCCCTCTCGCTC-3’) and the other in the piRNA binding region (GAATGTGACGGTAGGACCTG)—or alongside a nonbinding control guide (5’-GCACTGCTACGATCTACACC-3’).

### RT-qPCR

RNA was extracted from batches of wild-type embryos dissected within the area pellucida using TRIzol and reverse transcribed using Superscript III reverse transcriptase with random hexamers according to the manufacturer’s suggestions. qPCR was performed on an Eppendorf Realplex using MyTaq mix, SYBR green and the respective primers averaging CT values for technical triplicates. Chiwi and Chili Ct values at different developmental stages were normalized to 18s rRNA (Delta Ct) and fold difference was calculated to Chiwi levels at HH4 to analyze relative expression.

### Small RNA-seq

The midbrain region of WT HH9-chicken embryo heads was dissected and RNA was purified using TRIzol. Two replicates of several pooled embryos each were collected. 2ug of total RNA was then run on a 15% denaturing polyacrylamide gel and small RNAs within a 19-~38nt range were isolated and gel extracted as described previously^53^. Half of each sample was then oxidized in borate buffer (5x solution pH 8.6; 150 mM borax, 150 mM boric acid) and 25mM sodium periodate for 25 minutes at 25°C, while the other half of the sample was incubated in buffer only. Samples were then ethanol precipitated and libraries were cloned using the NEBNext Small RNA Library Prep Set for Illumina (E7330S) according to the protocol with NEBNext Multiplex Oligos for Illumina (set 1 E7335S). To optimize for the extremely low concentration of RNA and avoid overamplification of adapter dimers, adapters were diluted 1:10 for cloning of the oxidized samples. Libraries were sequenced on an Illumina Hiseq X Ten (150 bp reads, single end) at a sequencing depth of ~20 million reads. Reads were trimmed and mapped to the chicken genome (galGal6) using Bowtie^54^ with no mismatches end to end and multimapping reads included (-V 0 -k 1 --best). Reads were analyzed for sequence length, gene type they mapped to and whether they mapped sense or antisense to a feature. Reads mapping to TEs were extracted and analyzed for 5’ to 5’ complementarity using a previously published script that we altered to include reads ranging from 19-34 nucleotides^55^, and the ping pong z-score was calculated as described previously^56^ by taking the difference of the value at position 10 and the mean of the background values (values of all positions but 10), divided by the standard deviation of the background values. TE mapping reads were then analyzed for mapping orientation by read length, and after deduplication of libraries, sequence logos were generated for the first 18nt of each sequence using Weblogo^57^. Due to the low number of sense mapping reads and overrepresentation of some sequences, collapsing of libraries was necessary to resolve the 10A bias. Reads were normalized to reads per million mapped reads (to the genome) unless otherwise stated.

To generate a heatmap of small RNA reads mapping to TEs, reads mapping to different TE families were counted with FeatureCounts^58^ using a RepeatMasker GTF. Reads were then normalized by reads per million mapped reads to the genome, and hierarchical gene clustering was performed using Cluster 3.0^59^. Normalized read counts were then adjusted by log_2_ and a heatmap was generated using Morpheus software^60^.

To analyze small RNAs mapping to the Ens1 and ERNI loci, reads were mapped to chicken TE consensus sequences from Repbase^61^ using Bowtie with three mismatches allowed and reporting all valid alignments (-V 3 -a --best --strata). Reads were separately mapped to the ERNI mRNA sequence (refseq NM_001080874.1) using the same parameters. Reads were normalized to reads per million reads that map to the genome.

### mRNA-seq

Cranial neural folds of two replicate batches of HH9-embryos electroporated at HH4 with CRISPR control constructs on the left side and Chiwi CRISPR constructs on the right side were dissected and RNA was isolated using the RNAqueous-Micro total RNA isolation kit (Ambion). Libraries were prepared and sequenced on an Illumina Hiseq 2500 by the Millard and Muriel Jacobs Genetics and Genomics Laboratory at Caltech at a depth of ~60 million reads (50 base pair reads, single end). Reads were trimmed for adapter sequences and mapped to the galGal6 genome using Bowtie2^62^, and reads mapping to genes and TEs were counted with FeatureCounts. Differential expression analysis was performed using DESeq2^63^.

To analyze Chiwi and Chili levels in sorted cranial and trunk neural crest cells, previously published raw data (NCBI BioProject# PRJNA497902)^37^ was obtained and mapped to galGal6 using Bowtie2.

Transcripts per million (TPM) counts were generated with TPMCalculator^64^ using the Ensembl galGal6 GTF (GRCg6a, INSDC Assembly GCA_000002315.5)^65^, with the coordinates for the predicted Refseq Chili locus (XM_025142807.1) added, as Chili is not currently annotated in Ensembl. Read coverage plots were generated using the UCSC Genome Browser^66^ and normalized to reads per million mapped reads. Ensembl tracks were used for gene models in the figures for all transcripts except Chili, which is not annotated in Ensembl, and PLD6, which is annotated slightly differently in Refseq than Ensembl, with the Refseq version more closely matching transcript sequences from other vertebrates, as well as our RNA-seq data.

### Immunofluorescence

All embryos were fixed for 20 minutes at room temperature in 4% paraformaldehyde, and subsequently blocked in 10% goat or donkey serum in PBST (PBS, 0.5% Tween-20) for two hours at room temperature. Both primary and secondary antibody incubations occurred at 4°C for two nights in 10% goat or donkey serum, with four one hour washes in PBST at room temperature after primary, and two 30 minute washes in PBST after secondary antibody incubation. After imaging, whole mount embryos were post-fixed in 4% paraformaldehyde overnight at 4°C prior to sectioning. Primary antibodies used: Mouse IgG1 anti-Pax7 (1:10), Developmental Studies Hybridoma Bank. Secondary antibodies used: Molecular Probes donkey or goat secondary antibody conjugated to Alexa Fluor 488, 568, or 647 (1:1000).

### In situ Hybridization Chain Reaction (HCR)

All HCR was performed with probes designed by Molecular Technologies and following the published V3 protocol^67^. 20-probe sets were used for all genes except Sox2, for which a 12-probe set was used. Prior to sectioning embryos were postfixed in 4% paraformaldehyde overnight at 4°C.

### Sectioning

Cryosectioning was performed at a thickness of 18μm on a Microm HM550 cryostat. Embryo preparation included fixation in 4% paraformaldehyde overnight at 4°C (either from live embryos to post-fix processed embryos), followed 15% sucrose overnight at 4°C and 7.5% gelatin overnight at 37°C prior to mounting in silicone molds and snap freezing in liquid nitrogen.

### Imaging and statistical analysis

All images were taken using a Zeiss AxioImager.M2 with an Apotome.2. Whole mount images of cranial neural crest stained for PAX7 were analyzed in Fiji^68^ by measuring the area of the migratory crest on the experimental (right) side of the embryo and dividing it by the area of the migratory neural crest on the left (control) side. Cell counts of PAX7 positive cells in sections of the cranial region were taken with the Analyze Particles feature in Fiji. Fluorescence intensity was measured from maximum intensity projections of Z-stack images by manually drawing regions of interest to measure average intensity and subtracting average intensity of background regions. Experimental values were then divided by control values from the same image. For fluorescent intensity and cell count quantification, three non-adjacent cranial sections were measured and averaged to create a representative value for each embryo. For all statistical analysis on images, a paired two-tailed Student *t* test was performed to compare two values (experimental and control) within single embryos.

---

**Supplementary Figure 1:**
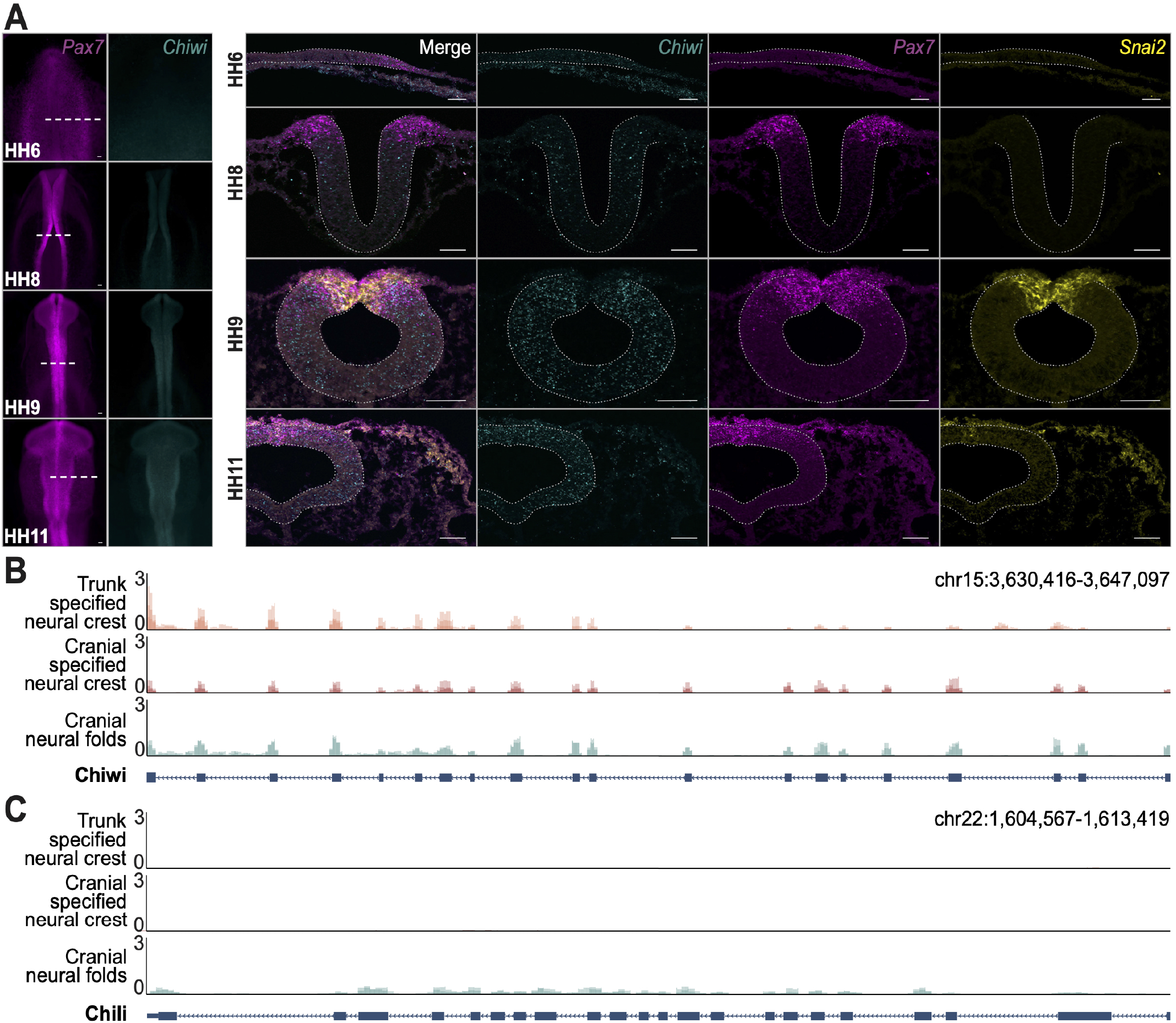
Chiwi and Chili expression in the neural crest. A) HCR depicts expression of *Chiwi* and neural crest markers *Pax7* and *Snai2* across a range of stages from HH6-HH12. Scale bars = 50μm. B-C) RNA-seq tracks of *Chiwi* (ENSGALT00000004171.6) and *Chili* (XM_025142807.1) expression in dissected cranial neural folds (n=2 biological replicates, control libraries from Fig 3A) as well as specified, FACS-sorted trunk and cranial neural crest cells (n=3 biological replicates; previously published data). Y-axis is reads per million reads mapped to genome. Replicate tracks are overlaid.

**Supplementary Figure 2:**
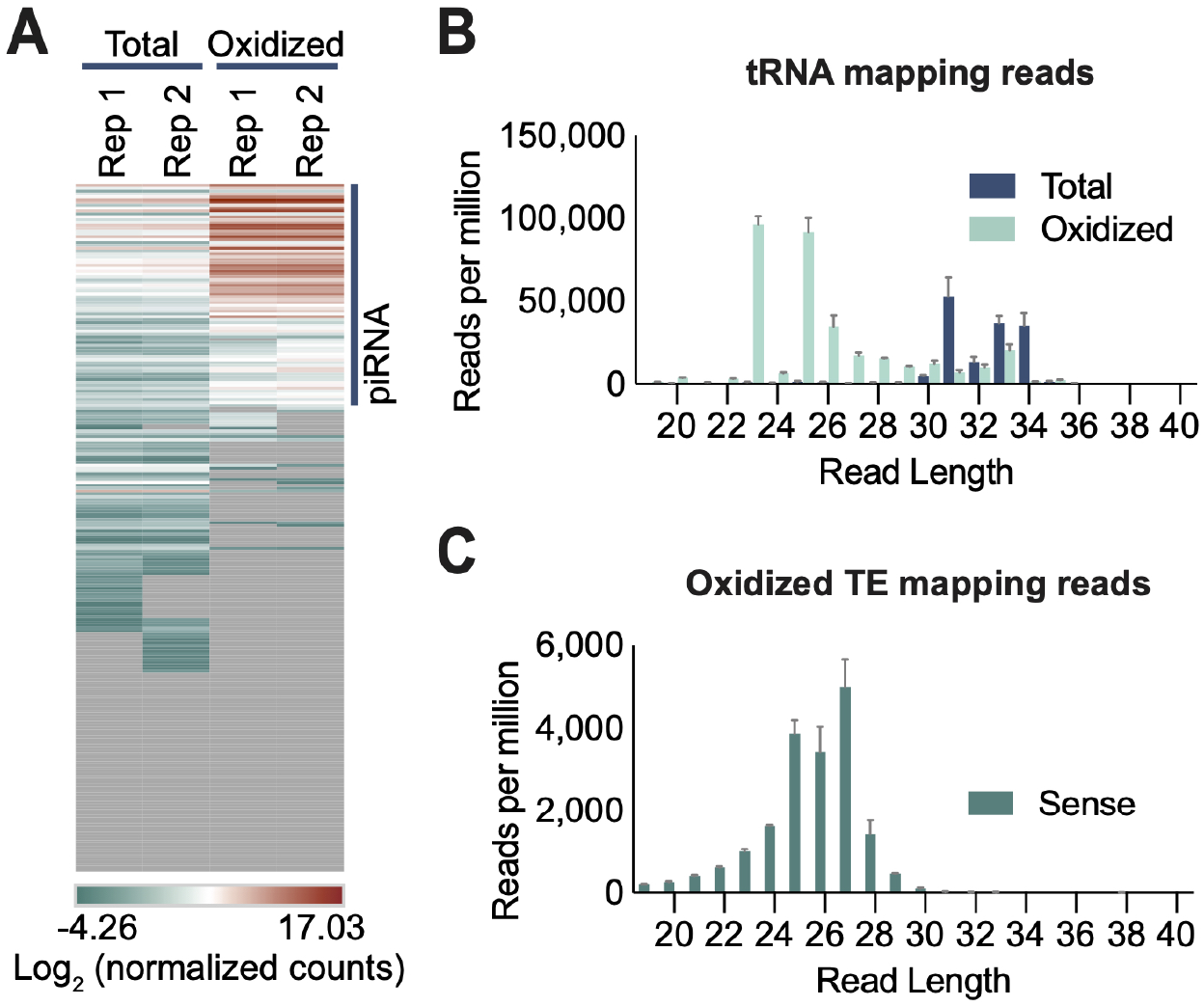
Small RNA sequencing extended data. A) Heatmap depicting normalized counts of small RNA sequences mapping to TEs. Each line represents a TE family. Counts are scaled by log_2_, after which the values of all cells range from −4.26 - 17.03. Cells with zero counts mapping are colored in gray (n=2 biological replicates). B) Length distribution of small RNA reads mapping to tRNAs (n=2 biological replicates). Error bars indicate st. dev. C) Length distribution of small RNA reads from oxidized libraries mapping to TEs in sense orientation (n=2 biological replicates). Error bars indicate st. dev.

**Supplementary Figure 3:**
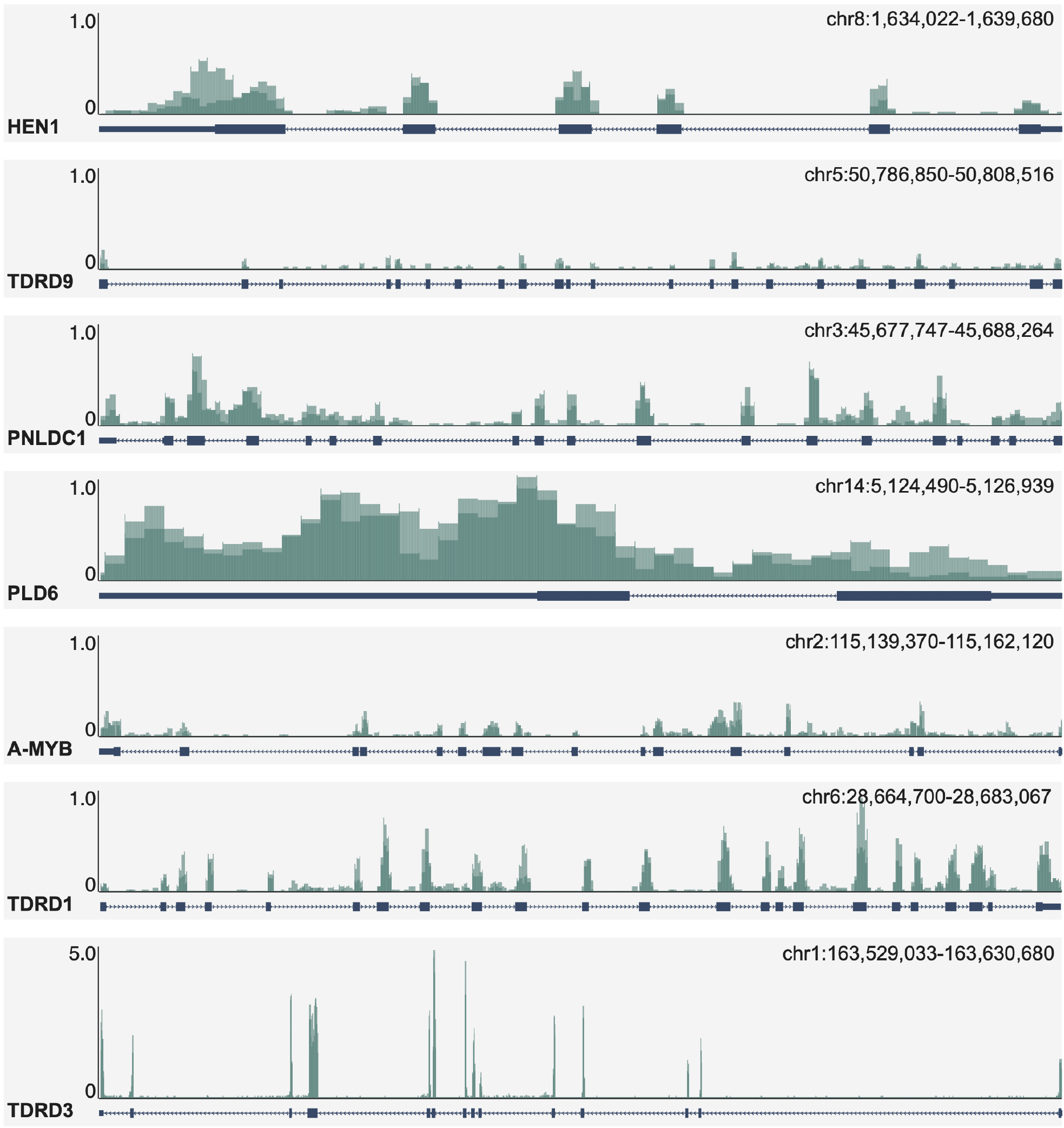
Expression of piRNA pathway genes in the neural folds. RNA-seq tracks of *Hen1* (ENSGALT00000003030.5), *TDRD9* (ENSGALT00000018863.7), *PNLDC1* (ENSGALT00000018991.6), *PLD6* (XM_015294407.2), *A-MYB* (ENSGALT00000066396.2), *TDRD1* (ENSGALT00000092521.1), and *TDRD3* (ENSGALT00000027376.6) expression in dissected cranial neural folds (n=2 biological replicates, control libraries from Fig 3A). Note that TDRD3 is shown on a larger scale than the other tracks. Y-axis is reads per million reads mapped to genome. Replicate tracks are overlaid.

**Supplementary Figure 4:**
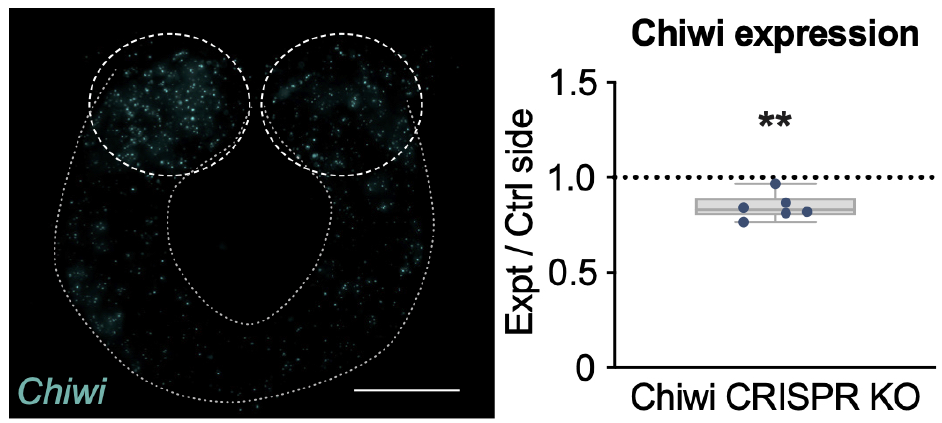
HCR confirmation of Chiwi CRISPR knockout. HCR confirms a reduction in *Chiwi* expression upon electroporation of CRISPR/Cas9 plasmids targeting Chiwi. Left: example of cranial cross sections. Scale bar = 50μm. Right: quantification of image analysis. Each data point represents the average fluorescent intensity of the right dorsal fold region (knockout, KO) divided by the left (control) side from three non-adjacent sections from the cranial region of a single embryo. Box plots indicate the interquartile range, while whiskers extend to min and max values. *, **, *** and **** indicate p values of ≤ 0.05, 0.01, 0.001, and 0.0001, respectively, and represent the difference between control and experimental meausrements.

**Supplementary Table 1:**
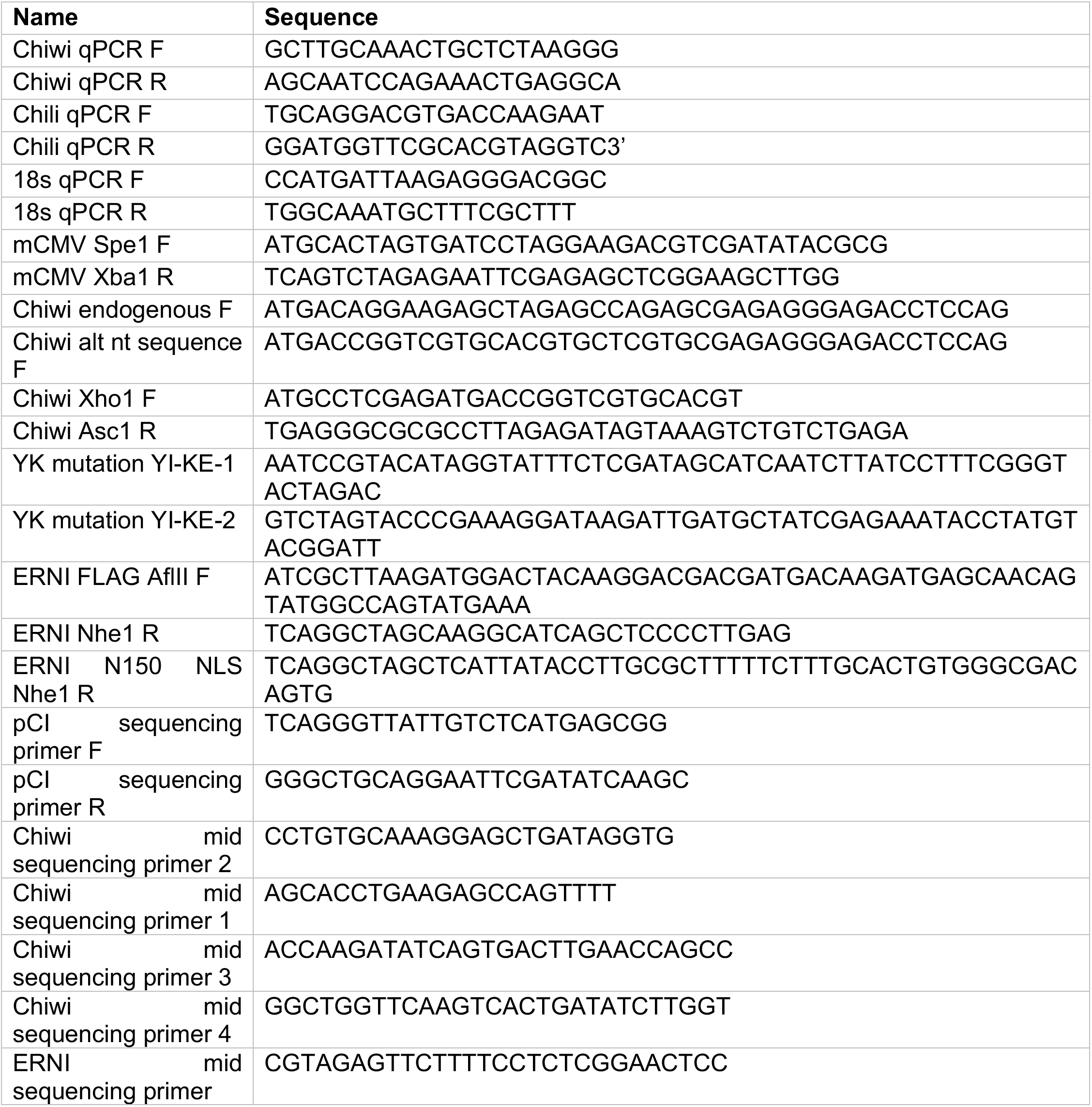
Primer sequences.

